# Sex, amitosis, and evolvability in the ciliate *Tetrahymena thermophila*

**DOI:** 10.1101/2021.12.21.473698

**Authors:** Jason Tarkington, Hao Zhang, Ricardo B. R. Azevedo, Rebecca A. Zufall

## Abstract

Understanding the mechanisms that generate genetic variation, and thus contribute to the process of adaptation, is a major goal of evolutionary biology. Mutation and genetic exchange have been well studied as mechanisms to generate genetic variation. However, there are additional processes that may also generate substantial genetic variation in some populations and the extent to which these variation generating mechanisms are themselves shaped by natural selection is still an open question. *Tetrahymena thermophila* is a ciliate with an unusual mechanism of nuclear division, called amitosis, which can generate genetic variation among the asexual descendants of a newly produced sexual progeny. We hypothesize that amitosis thus increases the evolvability of newly produced sexual progeny relative to species that undergo mitosis. To test this hypothesis, we used experimental evolution and simulations to compare the rate of adaptation in *T. thermophila* populations founded by a single sexual progeny to parental populations that had not had sex in many generations. The populations founded by a sexual progeny adapted more quickly than parental populations in both laboratory populations and simulated populations. This suggests that the additional genetic variation generated by amitosis of a heterozygote can increase the rate of adaptation following sex and may help explain the evolutionary success of the unusual genetic architecture of *Tetrahymena* and ciliates more generally.

## Introduction

Fisher’s fundamental theorem of natural selection states that the rate at which a population increases in mean fitness as a result of the operation of natural selection “is equal to its genetic variance in fitness at that time” (Fisher 1930). Genetic variants arise normally in populations through mutation, gene flow, and recombination and sex. In fact, it is hypothesized that sex is maintained, despite its costs, to generate genetic variance for fitness and increase evolvability (Weismann 1890; Kondrashov 1993; Burt 2000; Colegrave 2002).

*Tetrahymena thermophila* is a facultatively sexual, free-living, single-celled eukaryote with an unusual genome architecture that allows for the generation of additional genetic variation that we hypothesize should increase its evolvability following sex and through subsequent rounds of asexual division (Doerder 2014; Zufall 2016; Zhang et al. 2019). This increased evolvability may also provide a novel explanation for the maintenance of this unusual genome architecture.

Like other ciliates, *Tetrahymena* contain two types of nuclei: a silent germline micronucleus (MIC) and a transcriptionally active somatic macronucleus (MAC) (Merriam and Bruns 1988). The MAC gets destroyed after sex and a new one gets created from a mitotic product of the new zygotic nucleus (Orias 1986). In the model ciliate *T. thermophila*, the MAC contains *n* ≈ 45 copies of each of its 181 chromosomes (except the rDNA containing chromosome, which is present in *n* ≈ 9000 copies) and divides by amitosis (Fig. 1; Orias and Flacks 1975; Eisen et al. 2006, Sheng et al. 2020). During amitosis the chromosomes do not line up and segregate as they do during mitotic metaphase and anaphase. Instead the content of the genome is divided apparently randomly between the daughter cells (Karrer 2012). However, while *T. thermophila* is able to maintain ∼45 copies of every chromosome due to an unknown mechanism of chromosomal copy number control no such mechanism exist to maintain allelic diversity (Woodard et al. 1972; Larson et al. 1991; Sheng et al. 2020). In the absence of selection to maintain allelic diversity, over time amitosis will result in all but one allele being lost entirely from the MAC at each locus until the whole genome, except for *de novo* mutations, is homozygous (Fig. 1; Sonneborn 1974). This process is known as phenotypic or allelic assortment and occurs separately for all 181 MAC chromosomes. Amitotic division is thus predicted to result in large amounts of combinatorial genetic variation during the vegetative growth of a single sexual heterozygous progeny. This increase in genetic variation is expected to increase the rate at which an amitotically dividing population adapts (Doerder 2014; Zhang et al. 2019).

**Figure 1.**
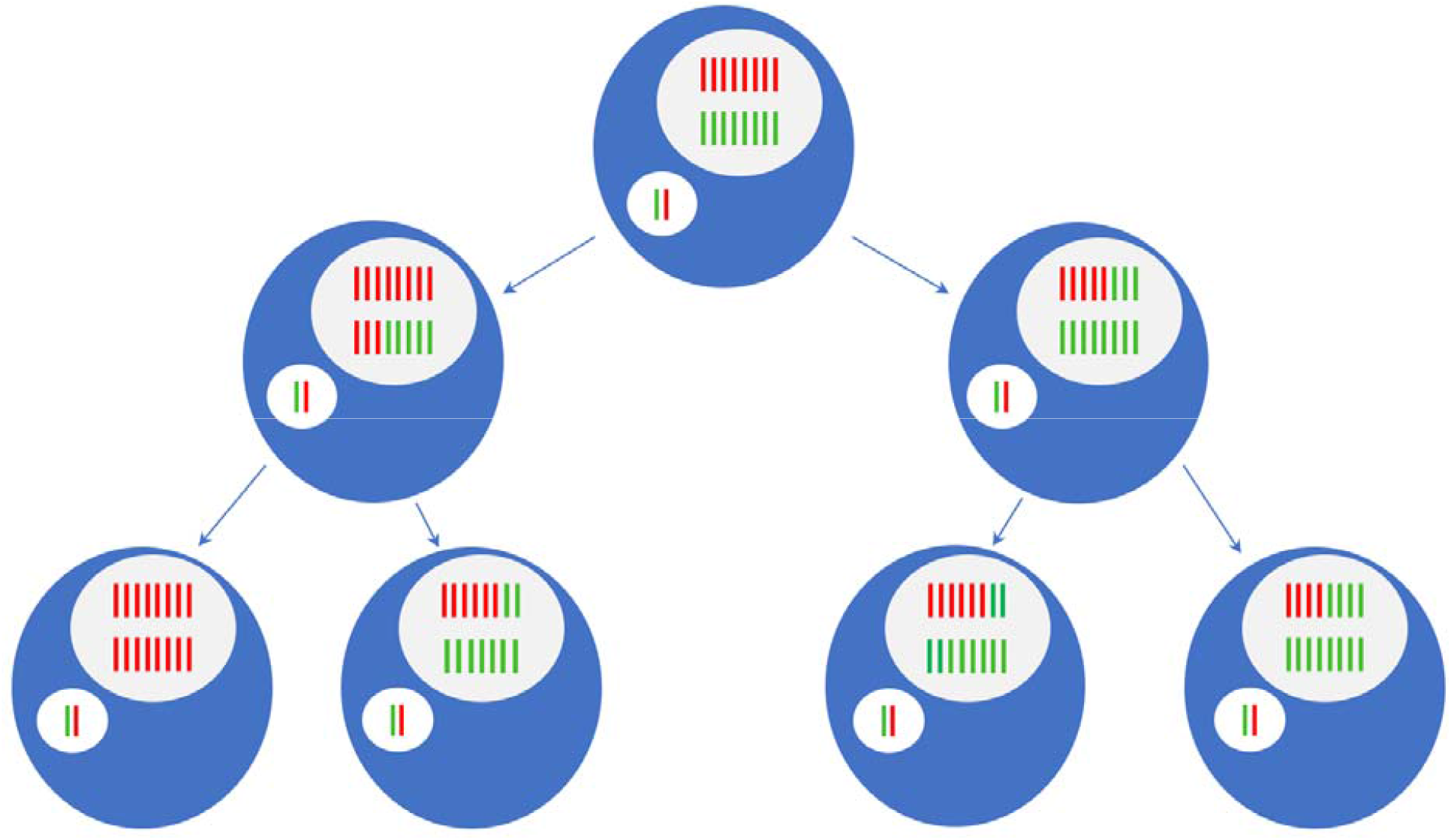
Amitosis and phenotypic assortment of a single chromosome. This figure shows the gradual loss of heterozygosity within a cell and increase in genetic variation among cells following sex. The small white oval is the diploid micronucleus (MIC) containing the red allele inherited from one parent and the green allele inherited from the other (only a single chromosome is shown for simplicity). The large white oval is the polyploid (*n*=16 here for simplicity) macronucleus (MAC). Following sex the macronucleus develops from the new zygotic nucleus and contains approximately half of the alleles from one parent and half from the other (shown in top cell). As the MAC divides amitotically each daughter cell (indicated by the arrows) inherits a random mixture of parental alleles. This process is accelerated in the figure above, while in reality it is likely to take ∼200 generations for 99% of loci to become homozygous (assuming that all alleles are neutral and a ploidy of *n*=45). Amitosis results in phenotypic assortment, i.e. the production of homozygous cells fixed for one or the other allele. Unlike mitosis, produces genetically variable progeny (as seen in the bottom row), which increases the genetic variation during the vegetative growth period following sex.

Previous models have shown that the unique genetic architecture of ciliates results in population genetics that differ from canonical population models (Morgens et al. 2014). Other models have shown that amitosis in *Tetrahymena* allows asexual lineages to slow Mueller’s ratchet and adapt at a rate similar to sexual lineages (Zhang et al. 2019). Additionally, several studies have claimed the genetic architecture of ciliates drives rapid gene and protein evolution (Zufall et al. 2006; Gao et al. 2014).

Typically, asexual progeny are genetically identical to their parents except for newly occurring mutations, but in *T. thermophila* a single heterozygous MAC genotype can give rise to a huge number of alternative genotypes among its asexual progeny. The high chromosome number in addition to recombination between homologous chromosomes (Deak and Doerder 1998) creates minimal physical linkage between loci allowing for differential assortment of most alleles. This results in many possible combinations of parental alleles being produced in a lineage descending from a single outcrossed individual. With selection acting on this variation, alleles and combinations of alleles will come to dominate in environments where they are advantageous, increasing the fitness of the population.

Here, we test the hypothesis that *T. thermophila* genome architecture and amitosis affect the dynamics of adaptation and the consequences of sex, specifically that populations founded from a single sexual progeny are more evolvable than populations founded from either unmated parent. To test whether amitosis will indeed increase the rate of adaptation following sex, we compare the rate of increase in replication time of progeny and parental populations during experimental evolution. We then use simulations to interpret and verify our experimental findings. The parental populations have already undergone phenotypic assortment and are thus expected to be largely homozygous in the MAC (Sonneborn 1974). This means that the effect of amitosis will be restricted to new mutations. In contrast, the sexually produced progeny should be highly heterozygous, meaning that amitosis will produce progeny with differing combinations of alleles. If amitosis increases evolvability following sex, then the rate of fitness increase in a population founded by a single new sexual progeny will be greater than that founded by an individual that has not had sex in many generations.

## Methods

### EXPERIMENTAL EVOLUTION

#### Overview

Three independent evolution experiments were performed to compare the rate of adaptation in populations derived from a sexual progeny to populations derived from individuals that had been dividing asexually only. Each experiment started with different parental genotypes of *T. thermophila*, which were crossed to produce a sexual progeny. A single cell was used to found the progeny populations. Populations were allowed to evolve for 1200-1500 generations, during which population growth rates were measured.

#### Strains and initial cross

For Experiment 1 we used natural isolates of *T. thermophila*, strains 19617-1 (which we refer to as B; *Tetrahymena* Stock Center ID SD03089; collected in Pennsylvania, USA; *cox1* GenBank: KY218380) and 19625-2 (A; collected in Pennsylvania, USA; *cox1* GenBank: KY218383; Doerder 2019). For Experiment 2 we used natural isolates 20453-1 (C; *Tetrahymena* Stock Center ID SD01561; collected in New Hampshire, USA; *cox1* GenBank: KY218424) and 20438-1 (D; ID SD01559; collected in New Hampshire, USA; *cox1* GenBank: KY218417). Experiment 3 used isolates 20395-1 (E; ID SD01557; collected in New Hampshire, USA; *cox1* GenBank: KY218412) and 20488-4 (F; ID SD01566; collected in Vermont, USA: *cox1* GenBank: KY218435).

Strains were thawed from frozen stocks, inoculated into 5.5 mL of the nutrient rich media SSP (Gorovsky et al. 1975) in a 50 mL conical tube, and incubated at 30 ºC with mixing for two days. These cultures were maintained as the parental lines. Eight (for Experiment 1) or 16 (for Experiments 2 and 3) populations were established for each genotype in 10 mL of SSP without shaking (for Experiment 1) or 180 µL of SSP with occasional shaking when the plates were on the microplate reader (for Experiments 2 and 3). For Experiment 1, four populations were maintained at 24 ºC and four at 37 ºC. For Experiments 2 and 3, all populations were maintained at 30 ºC.

To generate the hybrid progeny from these strains, a conical tube of each parental genotype was centrifuged and the supernatant was poured off before the cells were re-suspended in 10 µM Tris buffer (Bruns and Brussard 1974). After mixing at 30 ºC in Tris for two days to starve the cells and induce sexual competence, 1 mL of each starved parental population and an additional 1 ml of 10 µM Tris buffer were added to one well in a six-well plate and placed back in the 30 ºC incubator. The next morning (∼12 hours later) the plate was checked for pairs and put back in the incubator for an additional 4 hours to allow progression of conjugation. Individual mating pairs were isolated under a microscope using a 2 µL-micropipette and placed in 180 µL of SSP in one well of a 96-well plate. The plate was then incubated for 48 hours after which time a single cell was isolated from each well and re-cultured into 180 µL of fresh SSP in a new well. After another 48 hours at 30 ºC four (for Experiment 1) or 16 (for Experiments 2 and 3) individual cells, i.e. clones, were isolated from one of the wells, into new wells with SSP, one for each of the replicate populations, and incubated at 30 ºC for 48 hours. For Experiment 1, each of the four 180 µL cultures was then split in two with each half being added to a separate 50 mL conical tube containing 10 mL of SSP, one designated for evolution at 37 ºC and the other at 24 ºC.

We confirmed that progeny were indeed the product of sexual reproduction by performing maturity tests. Following successful sexual conjugation, *Tetrahymena* experience a period of immaturity when they will not pair or have sex again for approximately 60-100 generations (Doerder et al. 1995). Immaturity tests confirmed that our isolates would not pair, indicating that they were the recent progeny of sexual reproduction.

For Experiment 1, this provided us with a total of 24 populations consisting of three genotypes, two parental and one hybrid, half of which were evolved at 24 ºC and half at 37 ºC with four replicate populations of each genotype per treatment. For Experiments 2 and 3, there were 48 populations per experiment, consisting of 16 populations of each parent and the hybrids, all maintained at a single temperature (30 ºC).

Prior to the start of the experiment, parental strains had been kept in lab in cultures containing only a single mating type for at least 200 generations meaning that they have not had sex in at least that long and should therefore be highly homozygous in their MACs due to phenotypic assortment.

#### Experimental evolution transfer regime

In Experiment 1 approximately 20,000 cells (∼90 µL) from each 37 ºC culture and 60,000 (∼1 mL) from each 24 ºC culture were transferred to 10 mL of fresh SSP daily (Tarkington and Zufall 2021). Transfer volumes were adjusted as needed to maintain the same starting culture density at each transfer. On average, the 37 ºC evolved populations achieved ∼6.8 generations per day and the 24 ºC populations achieved ∼3.5 generations per day. We estimate the effective population size to be ∼70,000 cells for the 37 ºC environment and ∼128,000 for the 24 ºC environment by calculating the harmonic mean of the population size at the start of each discrete generation (Karlin 1968). Populations were maintained for over 4000 generations (Tarkington and Zufall 2021), but we only analyze the first 1500 generations here since this is the time during which amitosis is expected to have the greatest differential effect on heterozygotes versus homozygotes.

In Experiments 2 and 3, ∼1200 cells (2.625 µL) from each culture were transferred to 180 µL of fresh SSP in a 96-well plate daily and incubated at 30 ºC. This resulted in a starting density of ∼6700 cells/mL, a final density of ∼425,000 cells*/*mL, and ∼6 generations each day. These populations were evolved for 1500 (Experiment 2) or 1200 (Experiment 3) generations on a 96-well plate with an estimated effective population size of ∼3700.

#### Growth rate measurements

For Experiment 1 growth rate was measured by inoculating ∼500 – 1000 cells into one well of a 96-well plate and measuring the optical density (OD) at 650 nm in a microplate reader every 5 minutes over the course of 24 – 48 hours for 37 ºC assays and 48 – 72 hours for 24 ºC assays (Tarkington and Zufall 2021). The maximum growth rate was then estimated for each well by fitting a linear regression to the steepest part of the growth curve, estimating the maximum doublings per hour (h^-1^) (Wang et al. 2012; Long et al. 2013). 3 – 4 replicates of all populations were measured on a plate at each time point and the mean growth rate per plate was used in our analysis.

For Experiments 2 and 3, populations were evolved in 96-well microplates and typically alternated every 24 hours between being incubated in a 30ºC incubator and being incubated at 30ºC on the microplate reader when growth rates are measured for each population as described in the previous paragraph. This resulted in approximately 1 growth rate estimate per population every 12 generations for over 1500 generations for Experiment 2 and nearly 1200 generations for Experiment 3.

#### Data analysis

For each of the three experiments described above, the results were analyzed separately by plotting the growth rates over time then comparing the fitted slope of the growth rate trajectory of the progeny populations to that of the parents. The slope of the growth rate trajectory (or the evolvability) of each genotype was estimated from the data using a linear model (*growth rate ∼ genotype* + *generations* + *genotype*generations)* to estimate growth rate. The genotype*generations term corresponds to the slope of the growth rate trajectory or the evolvability. This approach provided us with a standard error of our estimate allowing us to assess whether the slopes or evolvabilities of different genotypes are significantly different from each other. For Experiment 1, which included larger population sizes than Experiments 2 and 3, the natural log of generation was used to linearize the data. This allowed us to fit the data using our linear model and then directly compare the genotype*generation term. A regression analysis found no correlation between the residuals and generations after transformation.

The absolute increase in growth rate (i.e., evolved -ancestral growth rate) was also calculated for each population by binning every 250 (Experiments 1 and 2) or 100 generations (Experiment 3). A pairwise Student’s *t*-test was used to test for significant differences in the total increase in growth rate between each genotype at each time point. This analysis allows us to determine the time period over which the progeny experience the greatest increase in evolvability.

### MODEL

#### Overview

We use the individual-based stochastic model described in Zhang et al. (2019) to simulate the evolutionary trajectory of *Tetrahymena* under different initial mutated allele distributions, particularly focusing on fitness and mutated allele fixation dynamics. Each individual carries *L* = 200 unlinked fitness loci within both the MAC and MIC. All mutations are assumed to be beneficial in the initial model, and act additively within a locus and multiplicatively among loci.

Two different initial mutated allele distributions are set to mimic the parent and progeny genotypes after sexual reproduction. For each parental genotype, *K* < *L* / 2 non-overlapping loci within the MAC are set to be homozygous for the beneficial allele, i.e., carrying *n* = 45 copies of a mutated allele initially; the remaining *L* – *K* loci are homozygous for a wildtype (unmutated) allele. The hybrid progeny have *2K* of the *L* loci in a heterozygous state, carrying either 22 or 23 mutated alleles. Hence, the parent and progeny genomes initially carry approximately the same number of beneficial alleles but differ in the way they are distributed throughout the genome. Consistent with Experiments 2 and 3, here we set the population size to *N* = 3,000 and number of replicates to 16.

#### Parameter exploration

We manipulate three parameters in our model: the genomic beneficial mutation rate *U*, the beneficial effect of a mutation *s*, and the number *K* of loci that are initially homozygous for the mutant allele within the parental genotype. Mutations are irreversible and, therefore, *U* represents the overall genomic mutation rate of a mutation-free genotype carrying *L* = 200 loci. We allowed *U* and *s* to vary between 0.01 and 0.1 in steps of 0.01. We considered seven values of *K*: 0, 5, 10, 15, 20, 25, and 30. Thus, we tried a total of 700 parameter combinations. For each combination, the simulation was repeated 3 times, and the mean results were then compared with the real fitness data obtained in Experiments 2 and 3.

The simulation and experimental data from Generation 0 to Generation 1,000 are first normalized by dividing by the ancestral fitness to calculate the relative fitness and then recorded every 50 generations for comparison with the experimental data (using the available generation numbers closest to 50, 100, 150, etc.). The simulation data were normalized by dividing by the fitness at generation 0. For the experimental data, a linear regression was performed for the first 200 generations, and the intercept was used as the estimate of the ancestral fitness value for normalization. For each data point, we calculated the sum of squared deviations (SSD) between simulation and experiment. Since there are 4 parental (C, D, E and F) and 2 progeny genotypes (CxD and ExF) in Experiments 2 and 3, we calculated an overall deviation metric as the sum of the average SSD for the parental populations and the average SSD for the progeny populations.

This overall deviation metric was computed for each of the 700 parameter combinations.

#### Allele fixation dynamics within populations

The parameter combination that achieved the greatest similarity between the results of simulations and experiments was then used to analyze the dynamics of fixation of beneficial mutations during evolution. Here we focus on the *2K* loci that shape the initial genetic difference between parent and progeny genotypes. Adopting these parameters into our stochastic model and raising the number of replicates to 100, we analyze the mean number of loci fixed for beneficial alleles and the mean number of beneficial alleles carried per individual within the *2K* loci for the parent and progeny population, respectively.

#### Deleterious mutations

Another set of *L* = 200 loci which can only accumulate deleterious mutations are integrated into the model described above to study the fitness dynamics in the presence of deleterious mutations. Based on results from Long et al (2013) and assuming the same mutation parameters within the MIC and MAC, we set the deleterious genome-wide mutation rate *U*_*d*_ = 0.21 within the macronucleus, with the additive effect of each copy of a mutated allele being –0.00244 and the total fitness effect *s*_*d*_ = –0.11 for a locus homozygous for the mutated allele. Other parameters are the same as above. Fitness dynamics are monitored for 1,000 generations and compared with the results that assume all mutations are beneficial.

### Results

#### Experimental Evolution

As is commonly seen in evolution experiments, all populations showed increases in growth rate over the course of evolution (Kryazhimskiy et al. 2014; Lenski et al. 2015). All populations in Experiment 1 showed greater increases in growth rate than in Experiments 2 and 3, possibly due to the larger population sizes in Experiment 1. As predicted, in all three experiments, the populations founded from a single heterozygous sexual progeny cell adapted more quickly than the homozygous parental populations (Fig. 2). The same result was seen at all three temperatures demonstrating that the increased evolvability of the progeny populations is not temperature dependent.

**Figure 2.**
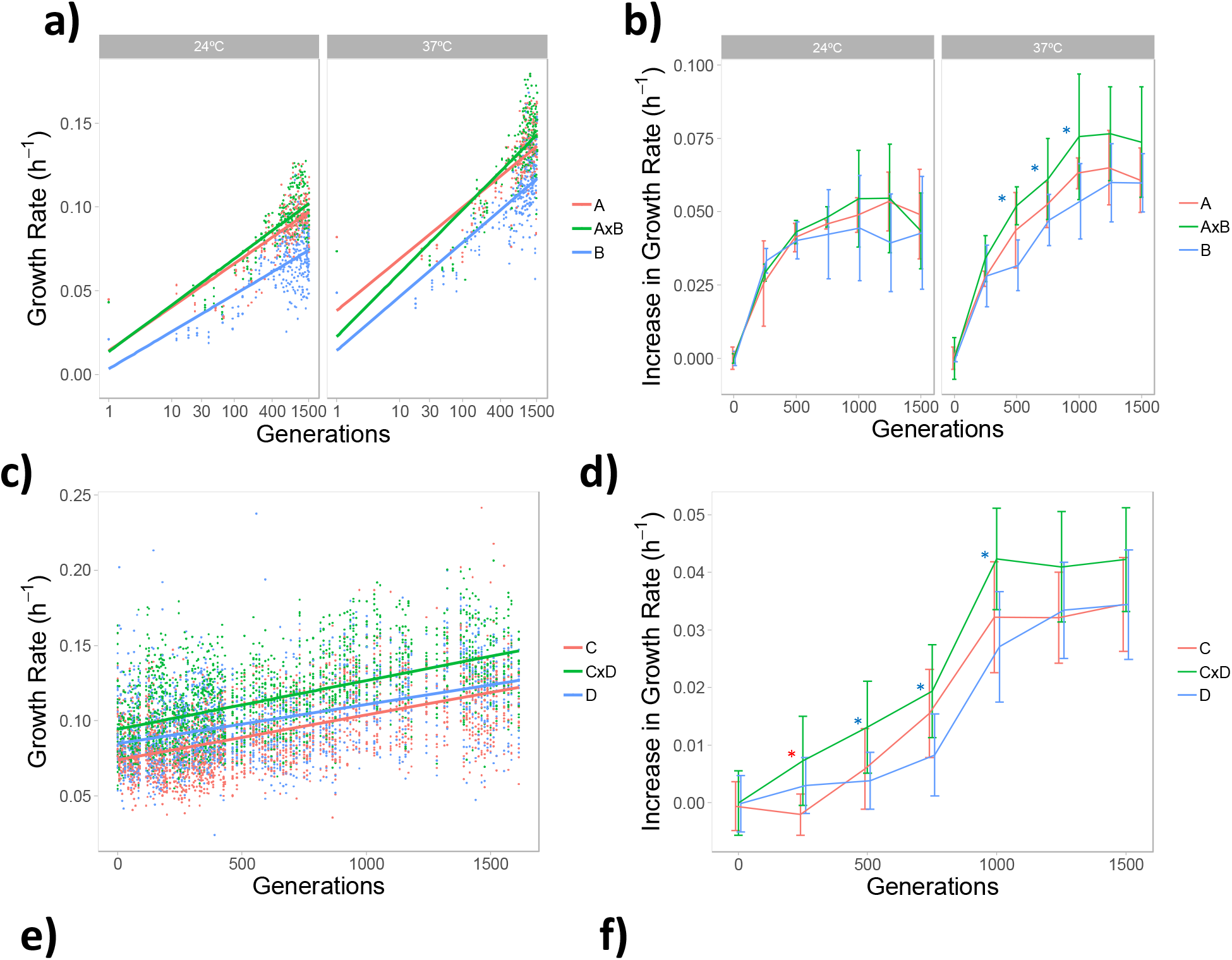
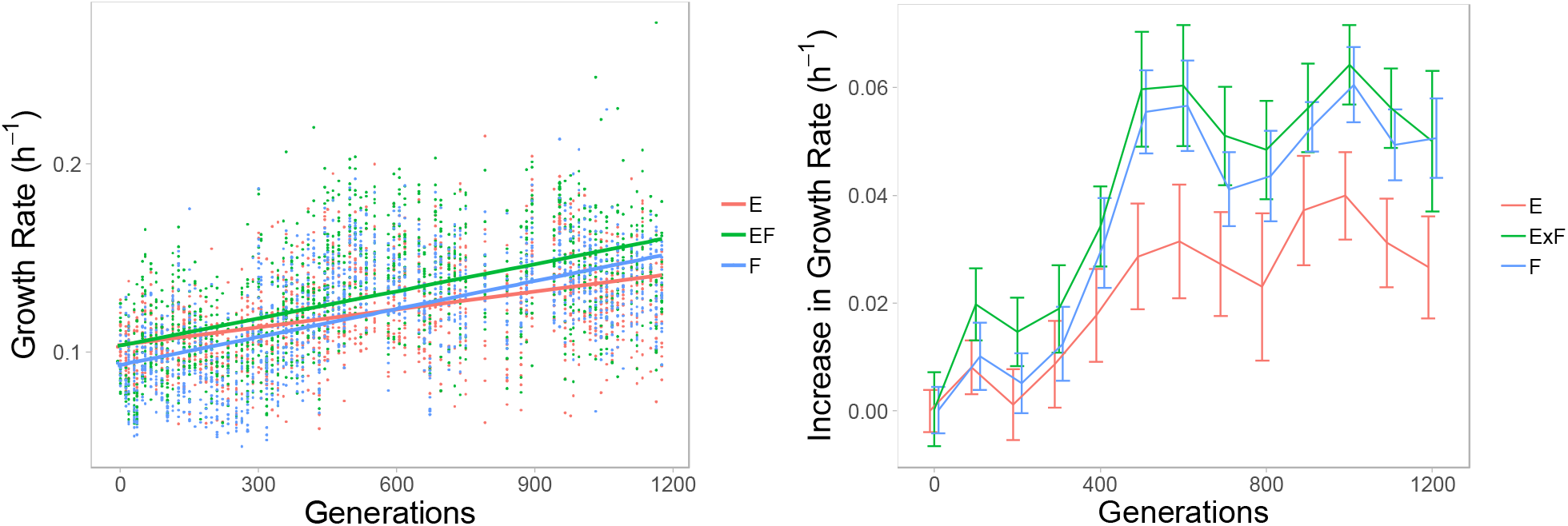
Growth rate trajectories of the parental (red and blue) and progeny (green) genotypes over 1200-1500 generations of evolution in Experiments 1 (a and b), 2 (c and d) and 3 (e and f). a, c, and e) Each point is the mean growth rate (panel a) or the growth rate (panels c and e) per time point of one of the replicate populations. A linear regression with a 95% confidence interval (shown in gray) is shown for each genotype in each experiment. b, d, and f). Lines connect the mean total increase in growth rate from the ancestor in each bin for each genotype. Error bars show the standard error of the 4 (panel b) or 16 (panels d and f) replicate populations in a 250 generation (panels b and d) or 100 generation (f) bin. The colored asterisks indicate the progeny has increased in growth rate significantly more than the parent of that color according to results of our Student’s *t* tests.

In Experiments 1 and 3 the initial fitness of the progeny was intermediate between the fitness of the two parents while in Experiment 2 the progeny initial fitness was higher than either parent. Despite the expectation from diminishing returns epistasis that genotypes with an initially lower fitness will increase in fitness faster (Kryazhimskiy et al. 2014; Wünsche et al. 2017), the progeny in Experiment 2 still increases faster in growth rate as expected due to increased genetic variation from amitosis.

In Experiment 1 we found that the progeny populations (AxB) increase in growth rate significantly faster than either parent (A and B) at both temperatures (Fig. 2a; estimate of slopes shown in Table 1). We also examined the total increase in growth rate from the ancestor (generation 0-125) every 250 generations and compared this between the progeny and parental populations. While the progeny populations increased more in growth rate on average at either temperature the small number of replicate populations (n=4) did not provide us with sufficient power to say whether this difference is significant for many time points especially at 24°C (Student’s *t*-tests shown in Fig. 2b).

**Table 1.** Estimates of evolvability. Estimates of the increase in growth rate per generation (or ln(generation) for Experiment 1) are shown in bold with standard error in parentheses. These estimates correspond to the slopes in figure 2 A, C, and E and are our measures of evolvability. The “*” indicates if the estimate of evolvability for parent 1 or 2 is significantly different than the estimate for the populations founded by a new sexual progeny. A standard least squares model including the effects of genotype, generation (ln(generations) for Experiment 1), and the interaction between them on r-max was used. Parent 1 column shows data for parents A, C, and E and Parent 2 for B, D, and F

|  | Sexual progeny x generation | Parent 1 x generation | Parent 2 x generation |
| --- | --- | --- | --- |
| Experiment 1 (24 °C) | **0.0178**<br>(0.00076) | **0.0147\***<br>(0.000744) | **0.0134\***<br>(0.000699) |
| Experiment 1 (37 °C) | **0.0237**<br>(0.000897) | **0.0190\***<br>(0.000867) | **0.0161\***<br>(0.000829) |
| Experiment 2 | **3.22x10<sup>-5</sup>**<br>(0.144x10<sup>-5</sup>) | **2.99 x10<sup>-5</sup> \***<br>(0.144 x10<sup>-5</sup>) | **2.60 x10<sup>-5</sup> \***<br>(0.144 x10<sup>-5</sup>) |
| Experiment 3 | **4.78x10<sup>-5</sup>**<br>(0.262x10<sup>-5</sup>) | **3.17x10<sup>-5</sup>\***<br>(0.263x10<sup>-5</sup>) | **4.97x10<sup>-5</sup>**<br>(0.262x10<sup>-5</sup>) |

In Experiment 2 the progeny populations (CxD) increased in growth rate faster than either of their respective parents (Fig. 2c; estimate of slopes shown in Table 1). The progeny populations (CxD) also had significantly greater total increases in growth rate than parent C after 250 generations and significantly greater increases than D after 500, 750, and 1000 generations (Fig. 2d). In Experiment 3 the progeny populations (ExF) increased in growth rate faster than parent E but not parent F (Fig. 2e; estimate of slopes shown in Table 1). However the progeny populations (ExF) had significantly greater absolute increases in growth rate than parent F after 100 and 200 generations and significantly greater increases than parent E at all time points (Fig. 2f).

#### Simulation

Among the 700 simulation parameter combinations, two of them, *U* = 0.03, *s* = 0.03, *K* = 5, and *U* = 0.10, *s* = 0.02, *K* = 10, generated fitness trajectories that most closely approximated the empirical ones (shown in Figure 3 among the 3 repeated simulations, the first set ranked 1^st^, i.e. most closely approximated the empirical trajectory, in one round and the second one ranked 1^st^ in another 2 rounds). The results do not differ qualitatively between these two combinations (comparison statistics SSD generated: 1^st^ round: 1.054 vs. 1.070, 2^nd^ round: 1.065 vs. 1.059, 3^rd^ round: 1.063 vs. 1.050), so below we only show the analyses based on the first parameter combination.

**Figure 3.**
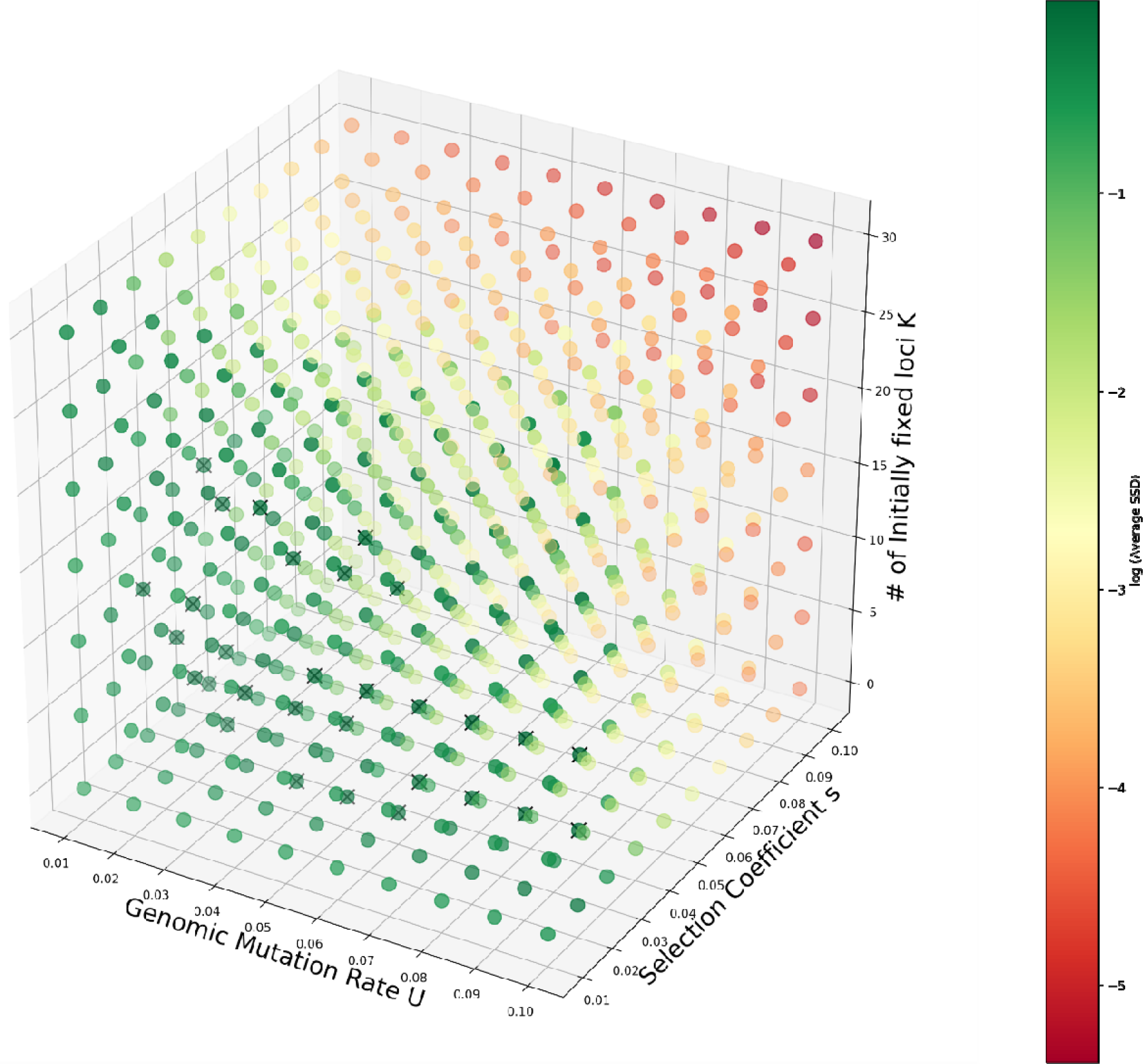
Certain simulation parameter sets achieve a better approximation to real empirical data from Experiments 2 and 3. Exploration of simulation parameters (genomic mutation rate *U*, selection coefficient *s* and the number of initially mutated allele fixed loci within the parent genotype *K*) to approximate the real empirical data. The color of the dots in the 3-dimension coordinate system represents the average sum of squared deviations (SSD) between simulation and experiment data. Both the genomic beneficial mutation rate *U* and selection coefficient *s* are set to vary between 0.01 and 0.1 in steps of 0.01; seven values of *K* ranging from 0 to 30 are tried. The parameters of the simulations are consistent with the real experiment, i.e., population size *N* = 3,000 and 16 replicates. Each parameter set was run in our stochastic model 3 times. The color bar on the right indicates the value of –log (average SSD), and the ones ranked as top 30 are marked as “x” in the graph.

The comparison of simulation and experimental results indicate that the observed evolutionary trajectories in the experiments may be explained by differences at ∼10 fitness loci between parents, a beneficial mutation rate of ∼0.03 per genome per generation and mutations increasing fitness by ∼3% in a homozygous state.

Consistent with our expectation, the progeny population exhibited a much faster fitness increase compared to the parental populations. During evolution over 1,000 generations, the population mean fitness (*ŵ*) of parental populations gradually increased from 1.16 to 1.61 on average, showing a relative fitness increase of 39% (Figure 4).

**Figure 4.**
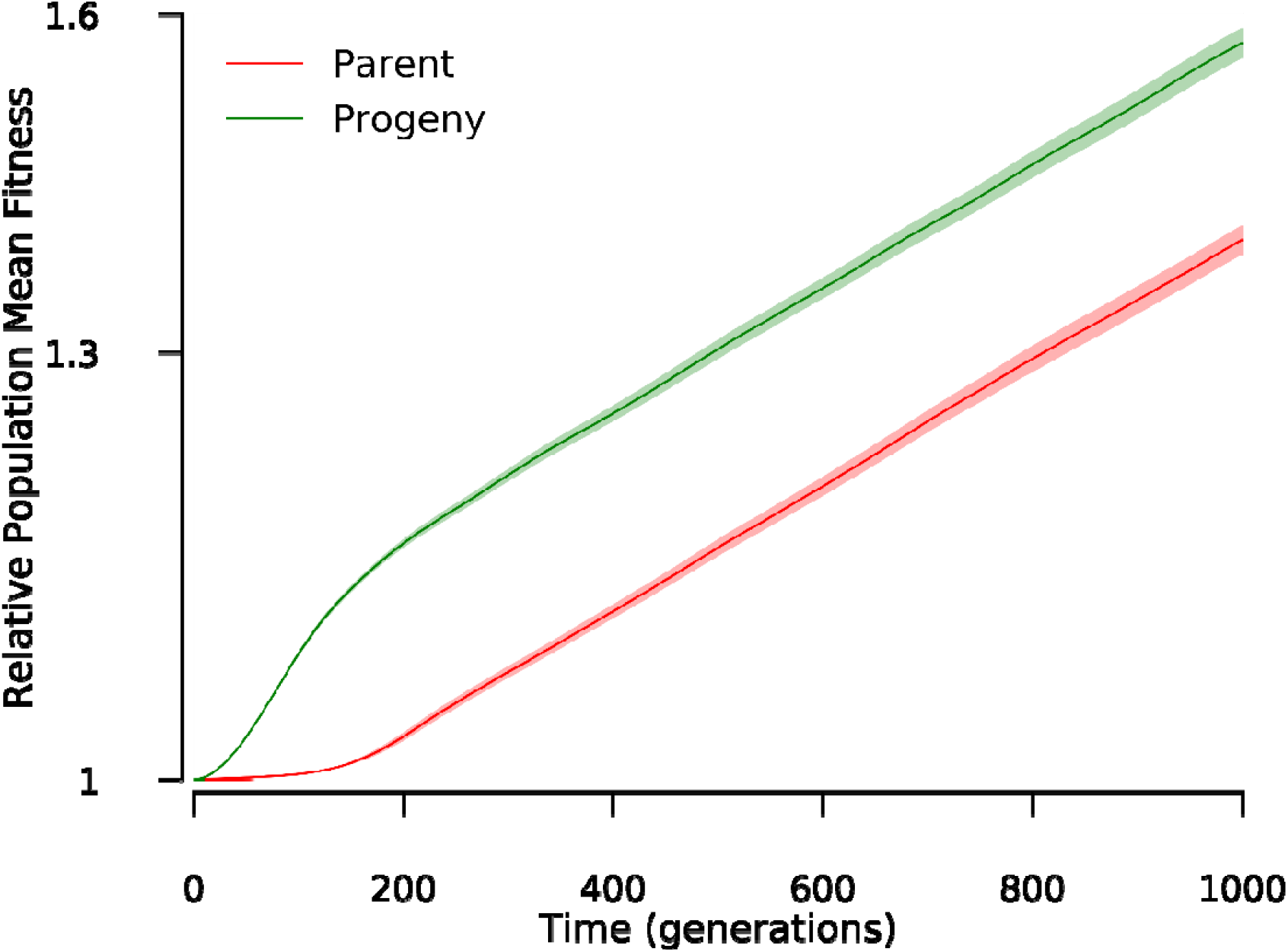
The progeny population exhibits a faster relative fitness increase compared to the parent population. The curves show the relative population mean fitness dynamics of the simulated parent and progeny populations during 1,000 generations of evolution. The actual fitness values from simulations are normalized by dividing by the initial population mean fitness to calculate relative fitness. The error bands represent 95% confidence intervals. Here we set genomic mutation rate *U* = 0.03, selection coefficient of beneficial mutations *s* = 0.03 and initially *K* = 5 loci are fixed in the parental genotypes. Note that the axis is shown in log scale.

However, with a similar starting fitness level of 1.16, the progeny population reached a much greater *w* of 1.82, a fitness increase of 57% (Figure 4). Using the slope between two consecutive generations as an indicator of the rate at which fitness increases, we found that the benefits of amitosis were mostly exhibited during the initial evolution process. Up to Generation 181, the progeny population showed a greater rate of fitness increase compared to the parent, and after that the two types of populations showed similar rates of fitness gain. The much faster fitness increase of the sexually reproduced progeny suggests that the evolvability of *Tetrahymena* is enhanced in the generations immediately following sex.

Using the simulation parameters explored above, we further explored fixation dynamics of beneficial alleles, which revealed distinct patterns between the parent and progeny populations. Here we only focused on the *2K* loci that shape the initial difference in allele composition between progeny and parent genotypes. Given *U* = 0.03, *s* = 0.03 and *K* = 5, an individual in the progeny population fixed an average of 9.87 (95% confidence interval: 9.80, 9.94) beneficial alleles among the *2K* = 10 loci at which they were initially heterozygous (Fig. 5A). In contrast, excluding the 5 initially fixed loci, only 0.19 (95% CI: 0.098, 0.282) of the remaining 5 loci become fixed for beneficial alleles within individuals in the parent population during the 1,000 generations of evolution.

**Figure 5.**
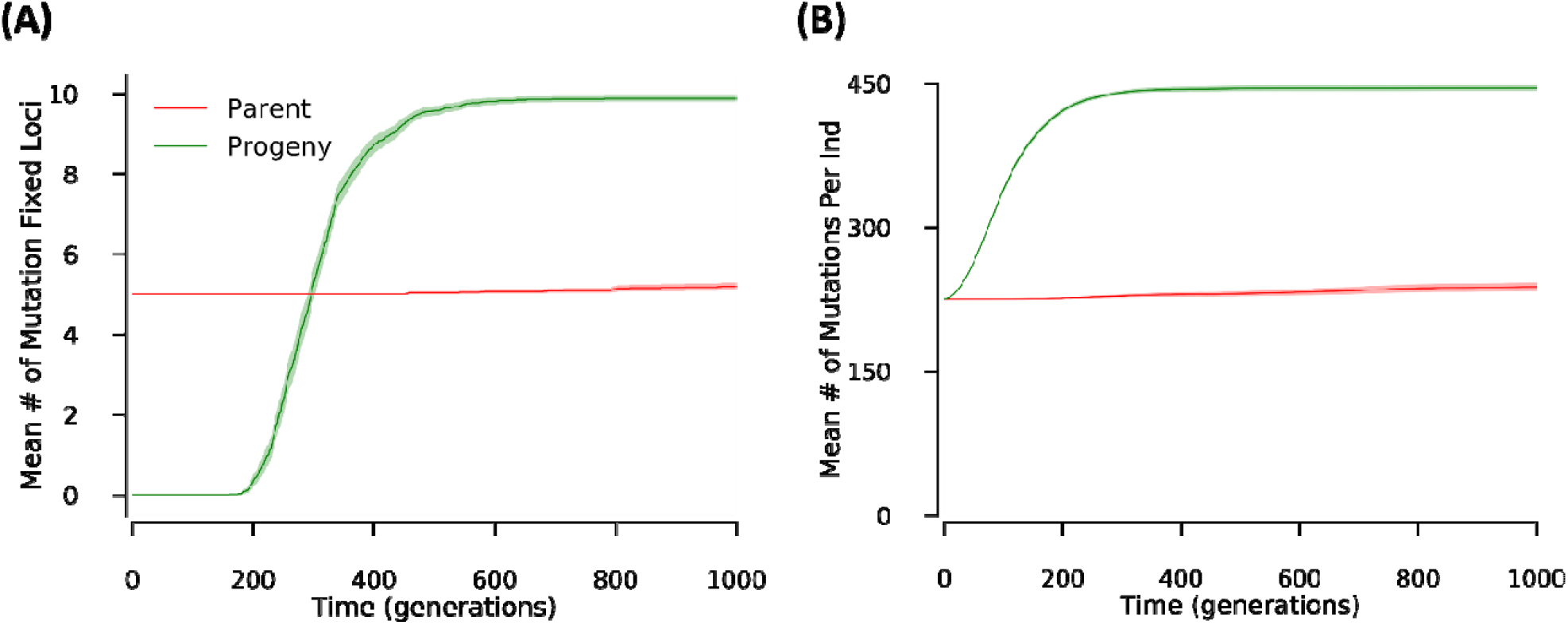
Parent and progeny populations exhibit distinct dynamics of fixation of beneficial alleles. The curves show the dynamics of (A) mean number of loci fixed for a beneficial allele and (B) mean number of beneficial alleles carried per individual for the *2K* loci at interest within the parent and progen population during the evolution of 1,000 generations. The simulation parameters are chosen from one of the 700 settings that match the experiment result best, i.e., *U* = 0.03, *s* = 0.03 and *K* = 5. Means and 95% confidence intervals are calculated from the simulation of 100 replicate populations.

Moreover, the beneficial allele fixation events within the progeny population are found to mainly occur between generation 200 and 400. From generation 0 to generation 200, although the total number of beneficial alleles gradually accumulated from ∼225 to ∼420 per individual, most of them are present as heterozygous (Figure 5B) with only ∼0.4 loci fixed for a beneficial mutation at generation 200 (Figure 5A). Then from generation 200 to 400, these beneficial alleles progressively reached fixation within the progeny population, as evidenced by the average mutations carried per individual being 443 (95% CI: 441, 446) and the mean number of mutation-fixed loci being 8.67 (95% CI: 8.46, 8.88) at generation 400. However, beneficial alleles accumulated very slowly in the parent population. The parent population only carried an average of 238 (95% CI: 233, 242) beneficial alleles per individual for the 10 loci at interest after evolving 1,000 generations.

Here we assumed that all mutations generated during evolution are beneficial, which is clearly unrealistic. However, our modified simulations demonstrate that the presence of deleterious mutations has little effect on the fitness dynamics. With a 7 times higher mutation rate and 3.7 times higher mutation effect compared to that of beneficial mutations, the model that incorporates deleterious mutations generated approximately the same fitness dynamics as the one assuming that all mutations are beneficial. After evolving for 1,000 generations and in the presence of deleterious mutations, the parent and progeny populations achieved relative mean fitness increases of 35% and 52%, respectively (Figure 6), which are very close to the values achieved when we ignored deleterious mutations. Using the linear regression slope between generations 500 and 1000 to measure the rate of fitness increase, both the parental and progeny populations did not differ significantly between two simulation regimes (two sample *t*-test, p = 0.13 and 0.82 for the parental and progeny population, respectively).

**Figure 6.**
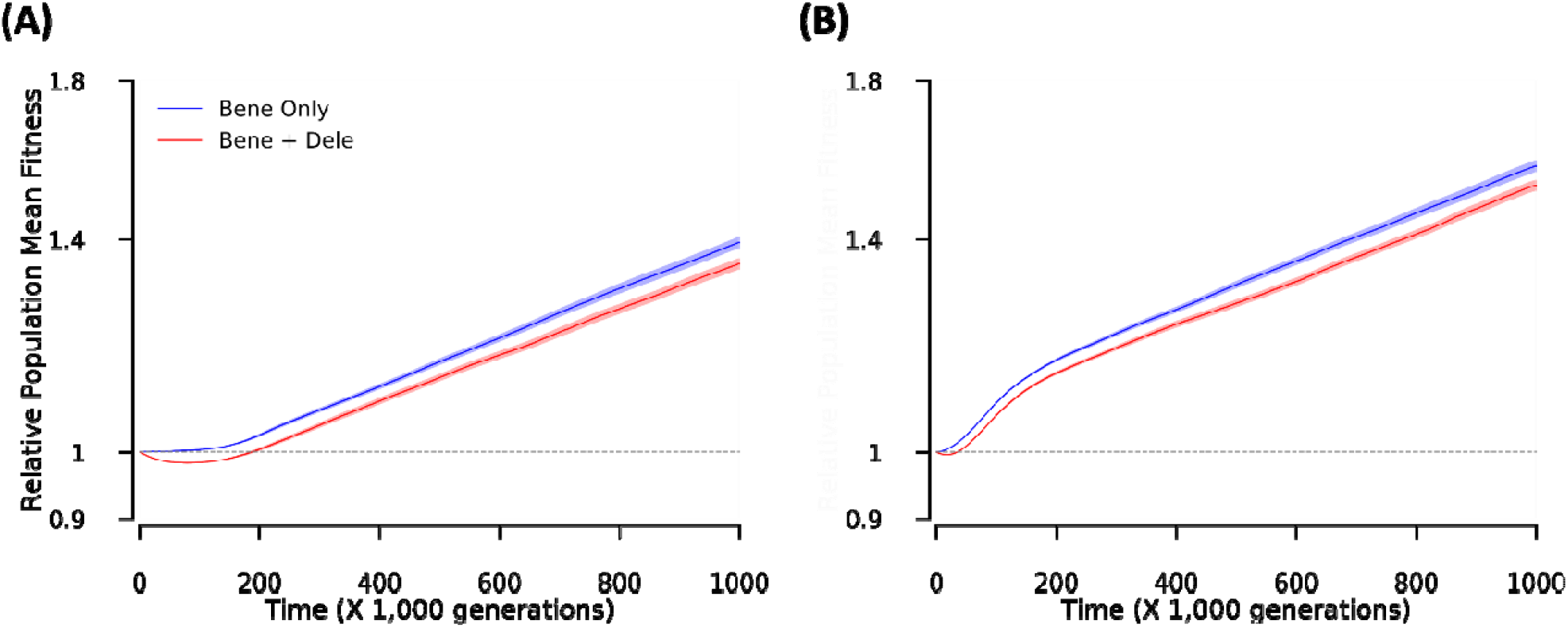
The presence of deleterious mutations has little effect on the fitness dynamics. The curves show the relative population mean fitness dynamics of the parent (A) and progeny populations (B) under the assumption of solely presence of beneficial mutations and co-presence of beneficial and deleterious mutations during the evolution of 1,000 generations. The actual fitness got in simulation are normalized by dividing the initial population mean fitness to calculate the relative fitness level. For the co-presence of beneficial and deleterious mutations, deleterious mutations are generated with a genomic mutation rate of *U*_*d*_ = 0.21 per generation, and decrease the fitness by *s*_*d*_ = –0.11 once present in homozygous state. Other parameters are the same as that in Figure 5. Note that the y axis is shown in log scale.

In summary, compared to the parental population, the progeny population exhibited a faster fitness increase, and distinct dynamics of fixation of beneficial alleles. Such a dramatic difference between the progeny and parental population results from the initial distribution of beneficial alleles within the genome. While dividing amitotically, the parental populations needs to wait for the generation of new mutations as there is no initial heterozygosity which can be segregated. However, the progeny population can immediately begin to segregate the beneficial alleles with the wild type alleles within each locus on which natural selection can then act.

## Discussion

We have demonstrated that populations of *T. thermophila* founded by a single newly produced sexual progeny are more evolvable than populations founded by their unmated parents. Because the parents are largely homozygous in their MAC due to many generations of asexual division, leading to phenotypic assortment (Merriam and Bruns 1988), and the progeny are generated from a cross between divergent strains and thus should be highly heterozygous in their MAC, a major difference between the parents and progeny is the degree of heterozygosity in their MACs at the start of the experiment. Thus, the increased evolvability that we observed in the progeny is likely attributable to amitosis generating additional population-level genetic variation in the lineages founded by a single heterozygous progeny. These findings have implications for our understanding of the evolution of ciliate and *Tetrahymena* genetic architecture and the evolution of evolvability more broadly. It also improves our understanding of the advantage of sex in *Tetrahymena* in addition to explaining the success of populations that are obligately asexual (Doerder 2014).

Using simulations based on our experimental design, we were able to estimate the number of loci, mutation rate, and fitness effects of mutations that could produce the results from our experimental populations. While it will be useful in the future to directly measure the proportion of beneficial mutations and the distribution of their effects in *T. thermophila*, it is interesting to compare our estimates to experimental estimates of these parameters from other species. In a barcoded yeast evolution experiment beneficial mutations with effect greater than 0.05 were estimated to occur at a rate of ∼1 × 10^−6^ per cell per generation and mutations with effect 0.02 < *s* < 0.05 at a rate of 5 × 10^−5^ (Levy et al. 2015). Another experiment using yeast estimated the beneficial mutations rate *s* = 0.01 and the rate *U* = 10^−4^ (Frenkel et al. 2014). Five estimates from *E. coli* have a median value for beneficial mutations of *s* = 0.03 (Hegreness et al. 2006; Lília et al. 2007; Sousa et al. 2012; Zhang et al. 2012), which is directly in line with the estimate from this study. In addition, our simulations highlight the importance of the first ∼200 generations following sex, thus future experiments should focus more finely on this period of increased evolvability.

It is also important to note some limitations of the model and experimental design. The model makes several assumptions about the underlying biology including the absence of dominance, overdominance, or underdominance. Additionally, there is no epistasis incorporated into the model and all mutations have a fixed and constant effect. For the experimental evolution we use growth rate as a proxy for fitness. While growth rate has been shown to be highly correlated with competitive fitness (Tarkington and Zufall 2021) it is possible that fitness increases in other portions of the growth cycle that are not fully captured in our growth curves.

The most widely accepted explanation for the unusual ciliate genome architecture is that genome duality evolved as a mechanism to allow foreign DNA to be sequestered in the germline (Bracht et al. 2013), and amitosis in the MAC is simply a consequence of the mechanism by which foreign DNA is eliminated from that genome. The fact that amitosis often leads to senescence and cell death in many ciliates (Simon and Orias 1987) was thus thought to be just an unfortunate side-effect of genome duality. However, we show that in *Tetrahymena* there is a period of increased evolvability following sex suggesting that amitosis can instead be beneficial, particularly in species with copy number control that prevents the loss of whole chromosomes during amitosis (Brunk and Navas 1992). This increased evolvability thus may have also contributed to the evolutionary success of the unusual genome architecture in ciliates.

Our results also have implications for the evolution of sex more broadly. Sexual reproduction is ubiquitous and ancient among eukaryotes and may be in part responsible for their massive diversification (Cavalier-Smith 2010). Despite this apparent dependence on sex (at least in the long-term) among eukaryotes, the nature of selection maintaining sex is not fully understood. Importantly, the selective benefits of sex must be quite strong to account for the various costs associated with sex (e.g., two-fold cost of sex, energetic costs of finding a mate, breaking up beneficial combinations of alleles; Gibson et al. 2017). One of the most robust theories for the success of sex is that sex provides an indirect benefit by increasing genetic variation in the population thereby allowing selection to operate more effectively to increase the population fitness (Weismann 1890; Kondrashov 1993; Burt 2000). This hypothesis can be contrasted with the direct benefits hypothesis in which sex increases the fitness of the parent or progeny directly (Kondrashov 1993). Indirect benefits have been demonstrated in several systems. For example, sex increases the rate of adaptation in populations of *Chlamydomonas* by increasing genetic variation among offspring (Colegrave 2002; Kaltz and Bell 2002). Direct benefits have also been shown in several systems. For example in facultatively sexual species such as the ciliate *Paramecium*, which must have sex to avoid senescence (Gilley and Blackburn 1994), sex provides a direct benefit. Here we show that in *Tetrahymena thermophila* a single sexually produced progeny has greater evolvability than either parent. This is a particularly interesting benefit of sex because although it is an indirect benefit as it takes many asexual generations and the action of selection for the benefit to manifest, it is unlike the indirect benefit of sex that Weismann spoke of which requires an entire population reproducing sexually (Weismann 1890). In *Tetrahymena* a single sexually-produced progeny results in increased genetic variation due to the amitotic asexual division following sex.

The increased evolvability following sex that we demonstrate here results from the dual nuclear architecture and amitosis that is specific to ciliates. However, amitosis of a heterozygous progeny may be considered analogous to a population that normally reproduces sexually by selfing but occasionally outcrosses (e.g. *Saccharomyces cerevisiae*). The outcross will generate heterozygosity in an individual progeny that can then subsequently be lost through loss of heterozygosity (LOH) events or through multiple generations of selfing. During these LOH events or rounds of selfing, combinatorial allelic variation will be generated among the descendants of a single outcrossing event. Thus, similar to the variation that is generated during rounds of amitosis following sex in *Tetrahymena*, a single episode of outcrossing among a selfing population could generate substantial genetic variation among the descendants potentially increasing their evolvability in much the same way that we observe for *Tetrahymena* (Morran et al. 2009). Likewise, LOH through gene conversion or other rare events resulting in uneven mitotic recombination in facultatively sexual species such as *Saccharomyces cerevisiae* may also provide a similar benefit to that which we observe in *Tetrahymena* (Smukowski Heil et al. 2017; James et al. 2019).

Despite these benefits of sex, *∼*50% of *T. thermophila* natural isolates are asexual. In fact, some of the oldest (∼10 million years) well-documented cases of asexual eukaryotes are *Tetrahymena* (Doerder 2014). The increased evolvability that we demonstrate here provides support for the hypothesis that amitosis is responsible for the success of these asexuals (Doerder 2014; Zhang et al. 2019). However, our results also suggest a reason that sexual reproduction is not lost entirely from *Tetrahymena. Tetrahymena* may be maximizing its capacity for adaptation via increased evolvability of the MAC, while minimizing the long-term risks associated with those adaptations, e.g. when the environment changes, by “resetting” the MAC following sexual reproduction (Orias 1986). This suggests that environmental change is likely to play a role in maintaining sex in ciliates in the long-run (Hinton and Nowlan 1987; Watson and Szathmáry 2016).

## References

Bracht, J. R., W. Fang, A. D. Goldman, E. Dolzhenko, E. M. Stein, and L. F. Landweber. 2013. Genomes on the edge: Programmed genome instability in ciliates. Cell 152:406–416.

Brunk, C. F., and P. A. Navas. 1992. Variable copy number of macronuclear DNA molecules in Tetrahymena. Dev. Genet. 13:111–117. John Wiley & Sons, Ltd.

Bruns, P. J., and T. B. Brussard. 1974. Pair formation in Tetrahymena pyriformis, an inducible developmental system. J. Exp. Zool. 337–344.

Burt, A. 2000. Perspective: sex, recombination, and the efficacy of selection—was weismann rightã Evolution (N. Y). 54:337–351. John Wiley & Sons, Ltd.

Cavalier-Smith, T. 2010. Origin of the cell nucleus, mitosis and sex: roles of intracellular coevolution. Biol. Direct 5:7.

Colegrave, N. 2002. Sex releases the speed limit on evolution. Nature 420:664–666.

Deak, J. C., and F. P. Doerder. 1998. High Frequency Intragenic Recombination During Macronuclear Development in Tetrahymena thermophila Restores the Wild-type SerH1 Gene. Genetics 148:1109 LP –1115.

Doerder, F. P. 2014. Abandoning sex: Multiple origins of asexuality in the ciliate Tetrahymena. BMC Evol. Biol. 14:1–13.

Doerder, F. P. 2019. Barcodes reveal 48 new species of tetrahymena, dexiostoma, and glaucoma: phylogeny, ecology, and biogeography of new and established species. J. Eukaryot. Microbiol. 182–208.

Doerder, P., M. A. Gates, F. P. Eberhardt, and M. Arslanyolu. 1995. High frequency of sex and equal frequencies of mating types in natural populations of the ciliate Tetrahymena thermophila. Proc. Natl. Acad. Sci. U. S. A. 92:8715–8718.

Eisen, J. A., R. S. Coyne, M. Wu, D. Wu, M. Thiagarajan, J. R. Wortman, J. H. Badger, Q. Ren, P. Amedeo, K. M. Jones, L. J. Tallon, A. L. Delcher, S. L. Salzberg, J. C. Silva, B. J. Haas, W. H. Majoros, M. Farzad, J. M. Carlton, R. K. Smith, J. Garg, R. E. Pearlman, K. M. Karrer, L. Sun, G. Manning, N. C. Elde, A. P. Turkewitz, D. J. Asai, D. E. Wilkes, Y. Wang, H. Cai, K. Collins, B. A. Stewart, S. R. Lee, K. Wilamowska, Z. Weinberg, W. L. Ruzzo, D. Wloga, J. Gaertig, J. Frankel, C. C. Tsao, M. A. Gorovsky, P. J. Keeling, R. F. Waller, N. J. Patron, J. M. Cherry, N. A. Stover, C. J. Krieger, C. Del Toro, H. F. Ryder, S. C. Williamson, R. A. Barbeau, E. P. Hamilton, and E. Orias. 2006. Macronuclear genome sequence of the ciliate Tetrahymena thermophila, a model eukaryote. PLoS Biol. 4:1620–1642.

Fisher, R. A. 1930. The genetical theory of natural selection. Clarendon Press. Oxford.

Frenkel, E. M., B. H. Good, and M. M. Desai. 2014. The Fates of Mutant Lineages and the Distribution of Fitness Effects of Beneficial Mutations in Laboratory Budding Yeast Populations. Genetics 196:1217 LP –1226.

Gao, F., W. Song, and L. A. Katz. 2014. Genome structure drives patterns of gene family evolution in ciliates, a case study using Chilodonella uncinata (protista, ciliophora, phyllopharyngea). Evolution (N. Y). 68:2287–2295.

Gibson, A. K., L. F. Delph, and C. M. Lively. 2017. The two-fold cost of sex: Experimental evidence from a natural system. Evol. Lett. 1:6–15.

Gilley, D., and E. H. Blackburn. 1994. Lack of telomere shortening during senescence in Paramecium. Proc. Natl. Acad. Sci. U. S. A. 91:1955–1958.

Gorovsky, M. A., M.-C. Yao, J. B. Keevert, and G. L. Pleger. 1975. Chapter 16 Isolation of Micro-and Macronuclei of Tetrahymena pyriformis. Methods Cell Biol. 9:311–327.

Hegreness, M., N. Shoresh, D. Hartl, and R. Kishony. 2006. An Equivalence Principle for the Incorporation of Favorable Mutations in Asexual Populations. Science (80-.). 311:1615–1617. American Association for the Advancement of Science.

Hinton, G. E., and S. J. Nowlan. 1987. How learning can guide evolution. Soc. Sci. Comput. Rev. 1:495–502.

James, T. Y., L. A. Michelotti, A. D. Glasco, R. A. Clemons, R. A. Powers, E. S. James, D. Rabern Simmons, F. Bai, and S. Ge. 2019. Adaptation by loss of heterozygosity in Saccharomyces cerevisiae clones under divergent selection. Genetics 213:665–683.

Kaltz, O., and G. Bell. 2002. The ecology and genetics of fitness in Chlamydomonas. XII. Repeated sexual episodes increase rates of adaptation to novel environments. Evolution (N. Y). 56:1743–1753.

Karlin, S. 1968. Rates of approach to homozygosity for finite stochastic models with variable population size. Am. Nat. 102:443–455.

Karrer, K. M. 2012. Nuclear dualism. Methods Cell Biol. 109:29–52.

Kondrashov, A. S. 1993. Classification of the hypothesis on the advantage of amphimixis. J. Hered. 84:372–387.

Kryazhimskiy, S., E. R. Jerison, and M. M. Desai. 2014. Global epistasis makes adaptation predictable despite sequence-level stochasticity. Science (80-.). 344.

Larson, D. D., A. R. Umthun, and Shaiu Wen-Ling. 1991. Copy Number Control in the Tetrahymena Macronuclear Genome. J. Protozool. 38:258–263. John Wiley & Sons, Ltd.

Lenski, R. E., M. J. Wiser, N. Ribeck, Z. D. Blount, J. R. Nahum, J. J. Morris, L. Zaman, C. B. Turner, B. D. Wade, R. Maddamsetti, A. R. Burmeister, E. J. Baird, J. Bundy, N. A. Grant, K. J. Card, M. Rowles, K. Weatherspoon, S. E. Papoulis, R. Sullivan, C. Clark, J. S. Mulka, and N. Hajela. 2015. Sustained fitness gains and variability in fitness trajectories in the long-term evolution experiment with Escherichia coli. Proc. R. Soc. B Biol. Sci. 282:20152292.

Levy, S. F., J. R. Blundell, S. Venkataram, D. A. Petrov, D. S. Fisher, and G. Sherlock. 2015. Quantitative evolutionary dynamics using high-resolution lineage tracking. Nature 519:181–186.

Lília, P., F. Lisete, M. Catarina, and G. Isabel. 2007. Adaptive Mutations in Bacteria: High Rate and Small Effects. Science (80-.). 317:813–815. American Association for the Advancement of Science.

Long, H.-A., P. Tiago, R. B. R. Azevedo, and R. A. Zufall. 2013. Accumulation of spontaneous mutations in the ciliate Tetrahymena thermophila. Genetics 195:527–540.

Merriam, E. V, and P. J. Bruns. 1988. Phenotypic assortment in Tetrahymena thermophila: Assortment kinetics of antibiotic-resistance markers, tsA, death, and the highly amplified rDNA locus. Genetics 389–395.

Morgens, D. W., T. C. Stutz, and A. R. O. Cavalcanti. 2014. Novel population genetics in ciliates due to life cycle and nuclear dimorphism. Mol. Biol. Evol. 31:2084–2093.

Morran, L. T., M. D. Parmenter, and P. C. Phillips. 2009. Mutation load and rapid adaptation favour outcrossing over self-fertilization. Nature 462:350–352. Nature Publishing Group.

Orias, E. 1986. Ciliate conjugation. Mol. Biol. ciliated protozoa 45–84. Academic Press San Diego.

Orias, E., and M. Flacks. 1975. Macronuclear genetics of Tetrahymena. I. Random distribution of macronuclear gene copies in T. pyriformis, syngen 1. Genetics 79:187–206.

Sheng, Y., L. Duan, T. Cheng, Y. Qiao, N. A. Stover, and S. Gao. 2020. The completed macronuclear genome of a model ciliate Tetrahymena thermophila and its application in genome scrambling and copy number analyses. Sci. China Life Sci., doi: 10.1007/s11427-020-1689-4.

Simon, E. M., and E. Orias. 1987. Genetic instability in the mating type system of Tetrahymena pigmentosa. Genetics 117:437–449.

Smukowski Heil, C. S., C. G. DeSevo, D. A. Pai, C. M. Tucker, M. L. Hoang, and M. J. Dunham. 2017. Loss of Heterozygosity Drives Adaptation in Hybrid Yeast. Mol. Biol. Evol. 34:1596–1612.

Sonneborn, T. M. 1974. Tetrahymena pyriformis. Pp. 433–467 in R. C. King, ed. Handbook of Genetics: Plants, Plant Viruses, and Protists. Springer US, Boston, MA.

Sousa, A., S. Magalhães, and I. Gordo. 2012. Cost of antibiotic resistance and the geometry of adaptation. Mol. Biol. Evol. 29:1417–1428. Oxford University Press.

Tarkington, J., and R. A. Zufall. 2021. Temperature affects the repeatability of evolution in the microbial eukaryote Tetrahymena thermophila. Ecol. Evol. n/a. John Wiley & Sons, Ltd.

Wang, Y., C. Diaz Arenas, D. M. Stoebel, and T. F. Cooper. 2012. Genetic background affects epistatic interactions between two beneficial mutations. Biol. Lett. 9:20120328–20120328.

Watson, R. A., and E. Szathmáry. 2016. How Can Evolution LearnãTrends Ecol. Evol. 31:147–157.

Weismann, A. 1890. Prof. Weismann’s theory of heredity. Nature 41:317–323.

Woodard, J., E. Kaneshiro, and M. A. Gorovsky. 1972. Cytochemical studies on the problem of macronuclear subnuclei in Tetrahymena. Genetics 70:251–260.

Wünsche, A., D. M. Dinh, R. S. Satterwhite, C. D. Arenas, D. M. Stoebel, and T. F. Cooper. 2017. Diminishing-returns epistasis decreases adaptability along an evolutionary trajectory. Nat. Ecol. Evol. 1:0061.

Zhang, H., J. A. West, R. A. Zufall, and R. B. R. Azevedo. 2019. Amitosis confers benefits of sex in the absence of sex to Tetrahymena. bioRxiv 794735.

Zhang, W., V. Sehgal, D. M. Dinh, R. B. R. Azevedo, T. F. Cooper, and R. Azencott. 2012. Estimation of the rate and effect of new beneficial mutations in asexual populations. Theor. Popul. Biol. 81:168–178.

Zufall, R. A. 2016. Mating Systems and Reproductive Strategies in Tetrahymena.Pp. 221–233 in G. Witzany and M. Nowacki, eds. Biocommunication of Ciliates. Springer International Publishing, Cham.

Zufall, R. A., C. L. McGrath, S. V. Muse, and L. A. Katz. 2006. Genome architecture drives protein evolution in ciliates. Mol. Biol. Evol. 23:1681–1687.

